## Supplemental Figures for "Behavioral discrimination and olfactory bulb encoding of odor plume intermittency"

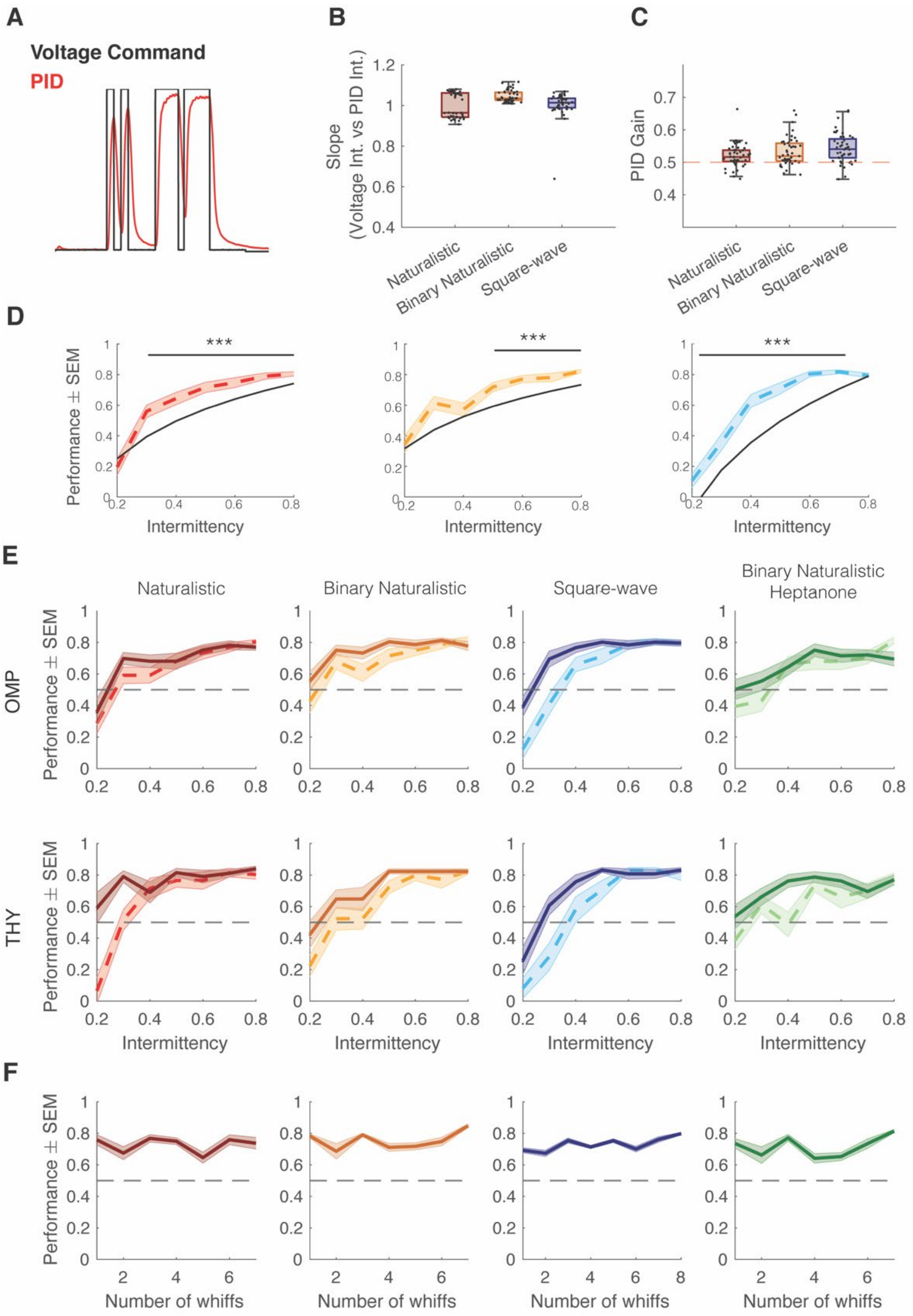


**Figure 1- figure supplement 1. Intermittency discrimination performance by genotype, odor, and whiff number.** (a) Example trial of voltage command and resulting PID reading. (b) Median slope of the correlation between voltage intermittency and PID intermittency for sessions of naturalistic (red), binary naturalistic (orange), and square-wave (blue) stimuli (n= 48 sessions per stimulus type, Naturalistic median = 0.96 IQR=[0.94 1.06] , Binary Naturalistic median= 1.04 IQR=[1.03 1.06] , Square-wave median= 1.01 IQR=[0.99 1.04]). (c) Median PID gain on 0.5 gain trials (n= 48 sessions per stimulus type, Naturalistic median = 0.52 IQR=[0.50 0.54] , Binary Naturalistic median= 0.52 IQR=[0.50 0.56] , Square-wave median= 0.54 IQR=[0.51 0.57]). One sample t-test with Ho:$\mu$ =0.5, p>0.05. (d) Performance on 0.5 gain trials (dotted colored line) and predicted performance of odor integration strategy at 0.5 gain (black line). One-tailed, one sample t-test with Bonferroni correction. Ho:$\mu_{Performance}$=Predicted Performance. Naturalistic intermittency≥0.3, p<0.0001; Naturalistic intermittency≥0.5, p<0.0001; Square-wave intermittency≤0.8, p<0.0001. (e) Performance curves by genotype (OMP-GCaMP6f and THY1-GCaMP6f). (f) Performance by number of whiffs in odor stimulus. Spearman correlation, p>0.05.

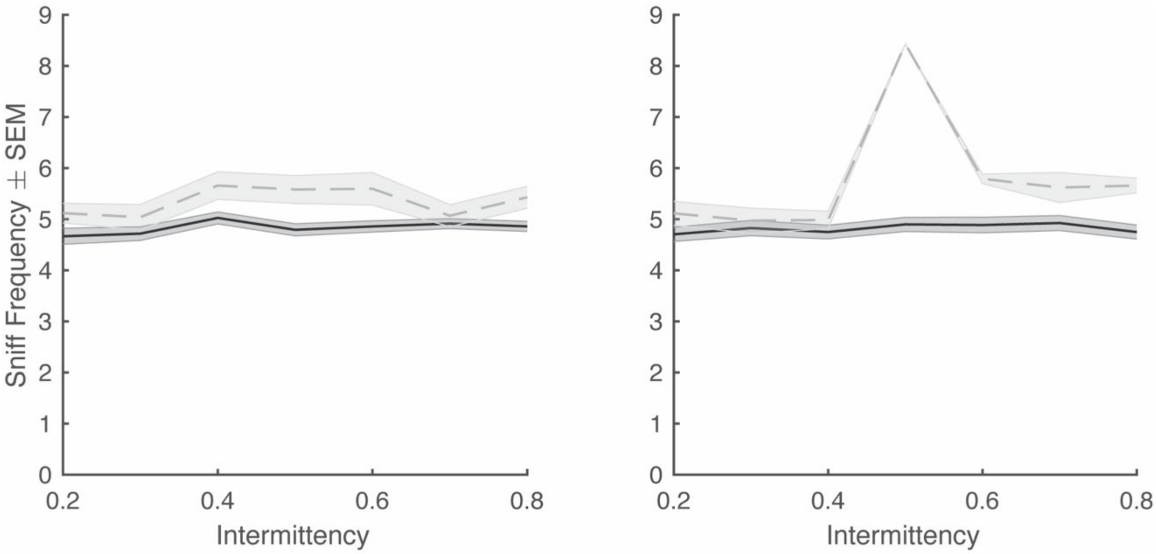


**Figure 2- figure supplement 1. Average trial sniff frequency vs odor intermittency on hit and miss trials.** Miss trials are in gray and hit trials are in black. *Left* Binary naturalistic. *Right* Synthetic (n=48 sessions each, generalized linear model; Sniff Frequency~ Trial Outcome*Intermittency, $p>0.05$; note: only one session with miss trials for synthetic stimuli at intermittency= 0.5).

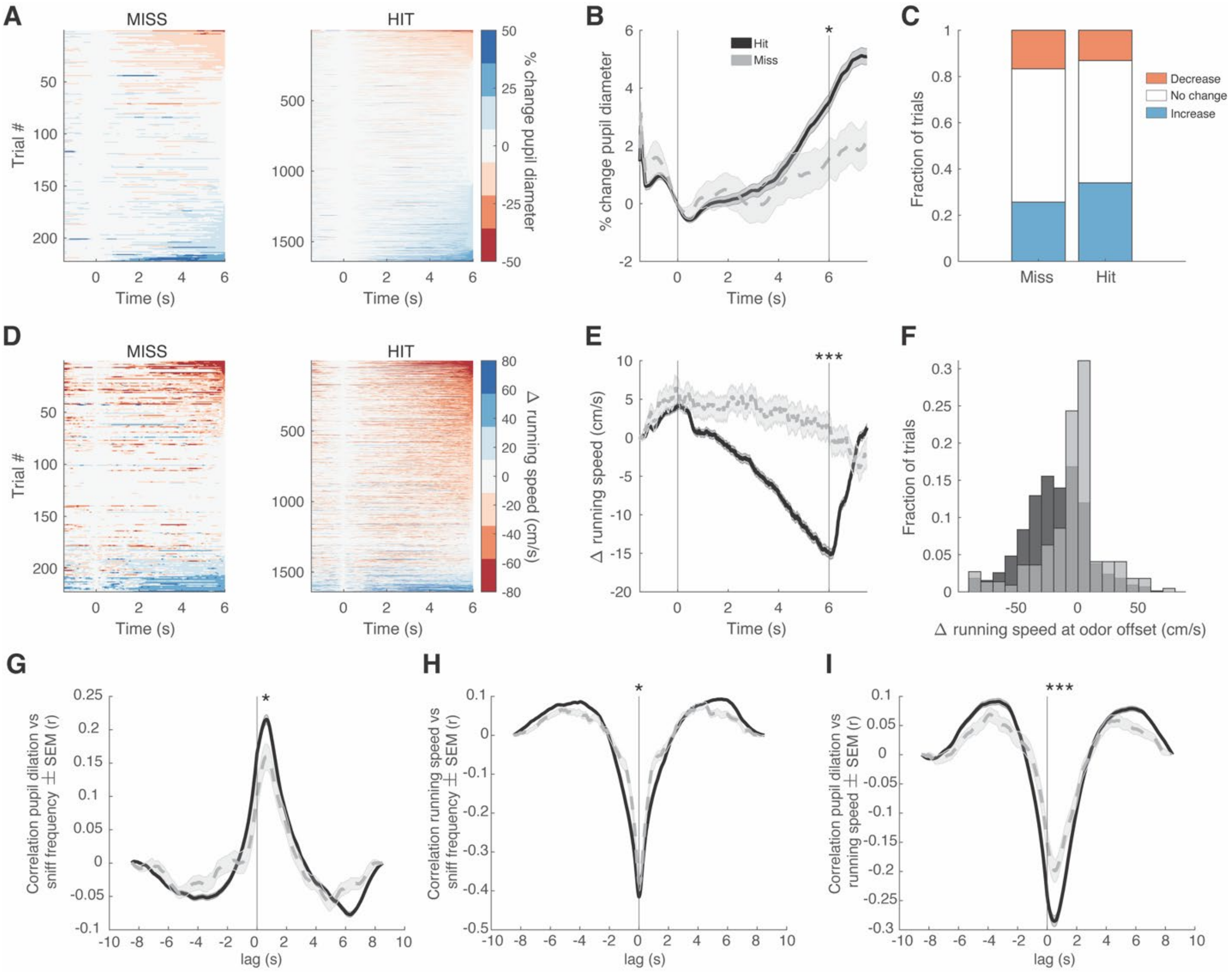


**Figure 2- figure supplement 2. Pupil dilation and running speed differ between hit and miss trials.** For all graphs miss trials are in gray and hit trials are in black. (a) Heatmap of the % change in pupil diameter from odor onset (time 0s) for miss and hit trials. (b) Average change in pupil diameter across trial time (odor onset is at time = 0s and odor offset is at time = 6s) for hit and miss trials. T-test at odor offset, $p<0.05$. (c) Fraction of trials with an increase, decrease, or no change in pupil dilation at the time of odor offset. Miss: increase, 25.7%; decrease, 16.7%; no change, 57.7%. Hit: increase, 34%; decrease, 13.1%; no change, 52.9%. (d) Heatmap of the change in running speed (cm/s) from odor onset (time 0s) for hit and miss trials. (e) Average change in running speed across trial time (odor onset is at time = 0s and odor offset is at time = 6s) for hit and miss trials. T-test at odor offset, $p<0.0001$. (f) Histogram of the change in running speed at odor offset for all trials. Two-sample Kolmogorov-Smirnov test, $p<0.0001$. (g) Cross-correlation of between pupil dilation and instantaneous sniff frequency during odor delivery period. Two-sample t-test at peak, $p<0.05$. (h) Cross-correlation between running speed and instantaneous sniff frequency during odor delivery period. Two-sample t-test at peak, $p<0.05$. (g) Cross-correlation between pupil dilation and running speed during odor delivery period. Two-sample t-test at peak, $p<0.0001$. In all cases a positive lag indicates a delay in the first parameter listed.

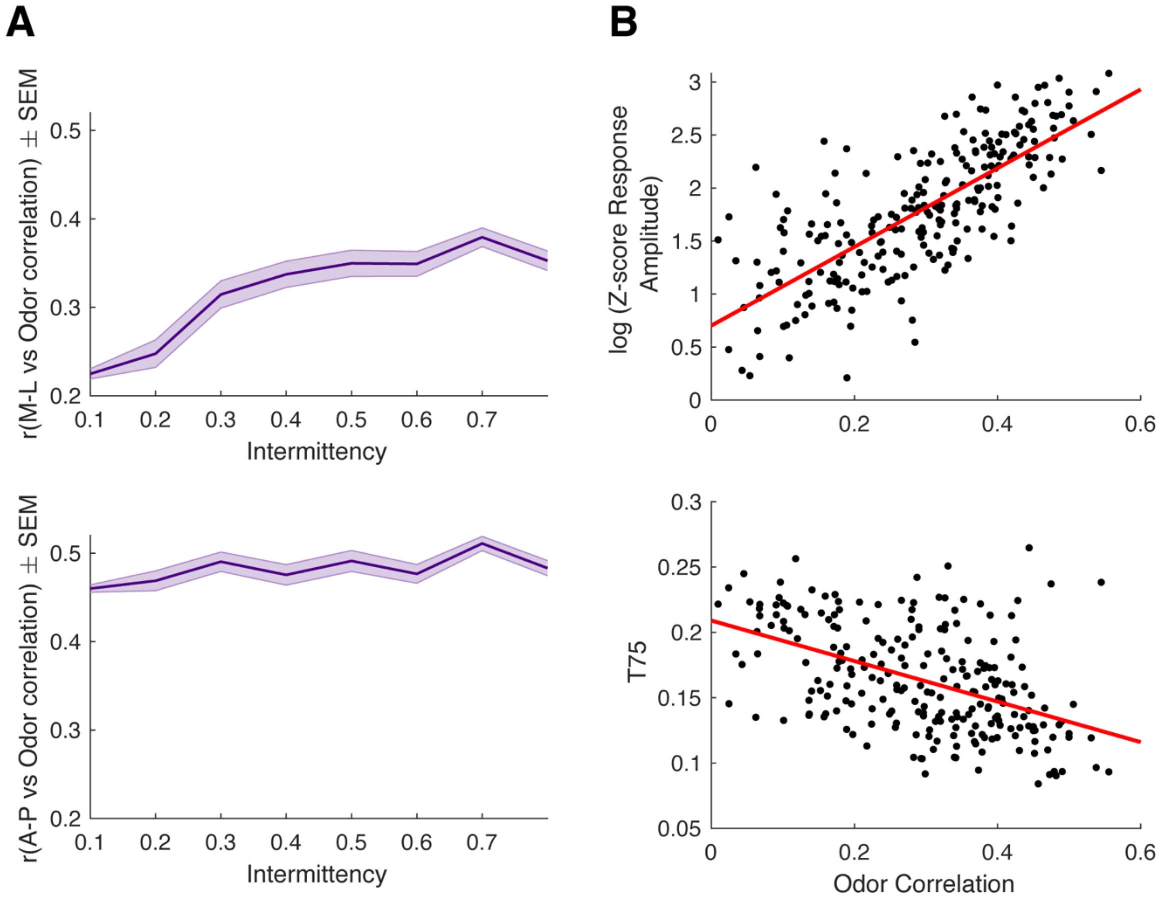


**Figure 3- figure supplement 1. Relationship between odor correlation and spatial location, response amplitude, and t75.** (a) Spatial correlation of odor-response correlation in M-L (*top*) and A-P (*bottom*) directions across intermittency values. M-L Linear regression: y=0.22x+0.21, p<0.0001, $r^2$= 0.08. A-P Linear regression: y=0.05x+0.46, p<0.0001, $r^2$= 0.009. (b) log(z-score response amplitude) vs odor correlation (y=3.71x+0.70, $r^2$=0.57, p<0.0001) and T75 vs odor correlation (y=-0.155x+0.21, $r^2$=0.23, p<0.0001).

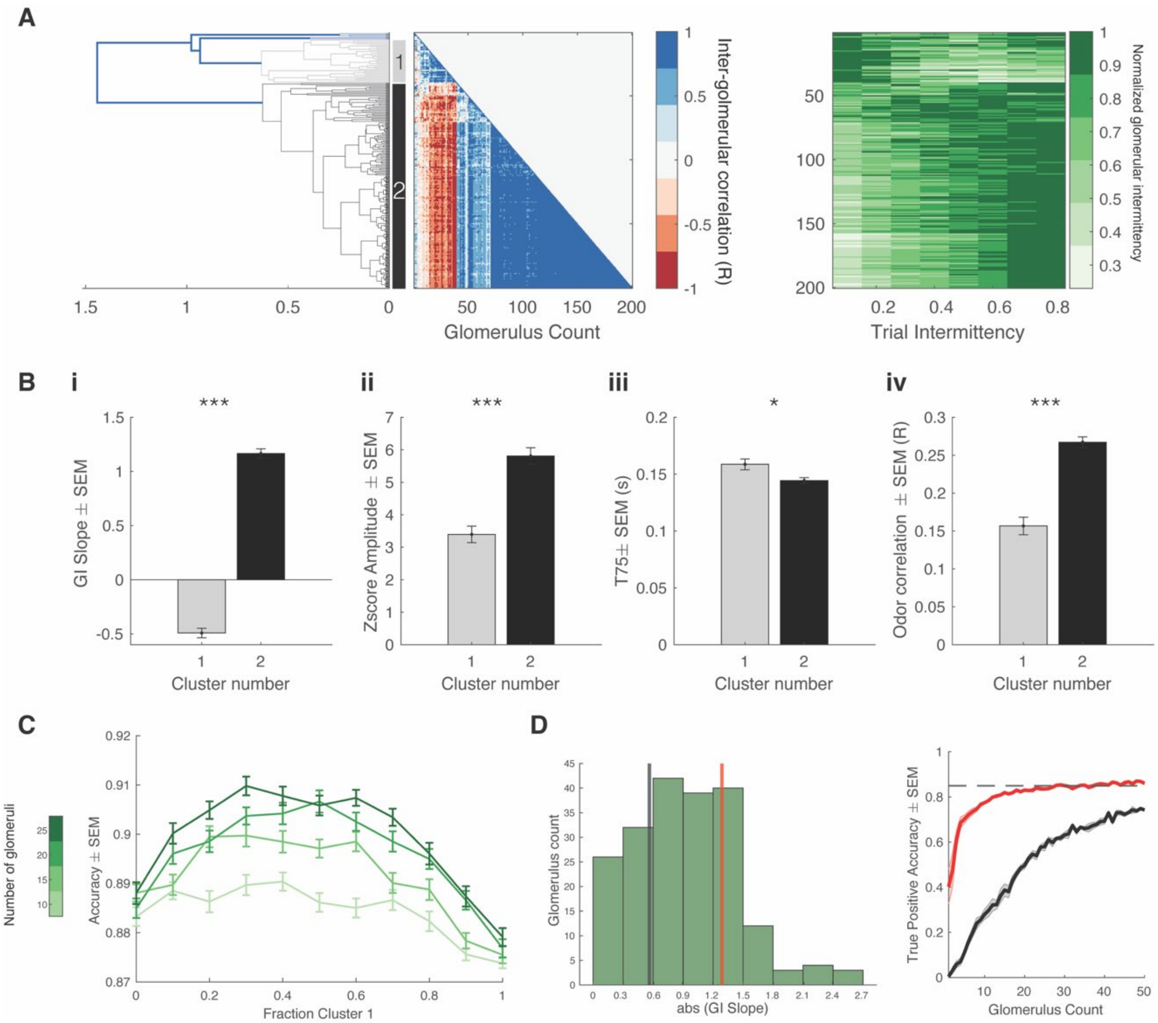


**Figure 4- figure supplement 1. Intermittency encoded in M/T cell glomerular subpopulations.** (a) *Left* Dendrogram for hierarchical cluster analysis. Gray indicates cluster 1 (34 glomeruli) and black indicates cluster 2 (161 glomeruli). *Middle* Inter-glomerular correlation matrix. Colorbar corresponds to correlation coefficient (r) between two glomeruli (glomerular intermittency vs odor intermittency). *Right* Color map of normalized glomerular intermittency. Rows sorted by hierarchical clustering. (b) i. Average slope of glomerular intermittency vs odor intermittency of cluster 1 and cluster 2 ($\mu_{cluster1}$=-0.49±0.04, $\mu_{cluster2}$=1.17±0.04) ii. Average z-score response amplitude for glomeruli in cluser 1 and cluster 2 ($\mu_{cluster1}$=3.4±0.25, $\mu_{cluster2}$=5.82±0.25) iii. Average T75 for glomeruli in cluster 1 and cluster 2 ($\mu_{cluster1}$=158.6±11.6 ms, $\mu_{cluster2}$=144.4±2.5 ms) iv. Average correlation between glomerular deconvolved ΔF/F traces and PID reading for glomeruli in cluster 1 and cluster 2 ($\mu_{cluster1}$=0.16±0.005, $\mu_{cluster2}$=0.27±0.007). (c) Accuracy of linear classifier trained using 10, 15, 20, and 25 glomeruli (colorbar) at varying fractions of cluster 1 and cluster 2 glomeruli. (d) *Left* Histogram of abs(GI Slope) for all glomeruli. Black line indicated the bottom 25$^{th}$ percentile (0.56) and red line indicates the top 25$^{th}$ percentile (1.29). *Right* True Positive Accuracy (CS+ predicted as CS+) of linear classifier trained on 0-50 glomeruli for glomeruli with the top 25$^{th}$ percentile of GI Slopes (red) and the bottom 25$^{th}$ percentile of GI Slopes. Dashed line indicates hit rate of animals on behavioral task (0.85).

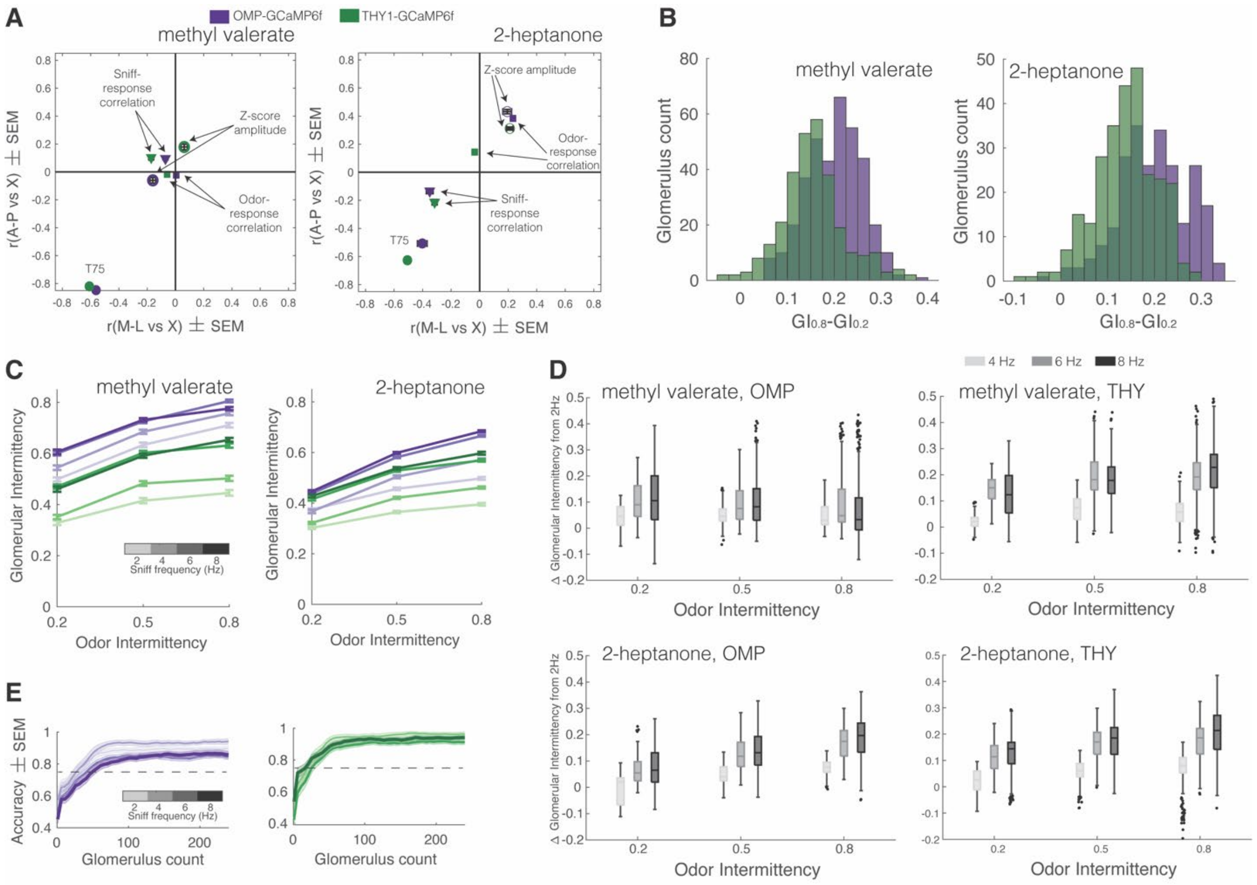


**Figure 5- figure supplement 1. Additional quantification of the effect of sniff frequency on glomerular representation of intermittency.** For all graphs purple indicates OSNs (OMP-GCaMP6f) and green indicates M/T cells (THY1-GCaMP6f). (a) Spatial maps of z-scored response amplitude (open circles) and odor-response correlation (diamonds) as well as spatiotemporal map (T75, filled circles) for methyl valerate (OMP, [M-L,A-P]: $\mu_{\text{z-score amplitude}}$=[-0.16,-0.06], $\mu_{\text{odor-response corr}}$=[-0.01,-0.02], $\mu_{T75r}$=[-0.56,-0.85]; THY, [M-L,A-P]: $\mu_{\text{z-score amplitude}}$=[0.06,0.17], $\mu_{\text{odor-response corr}}$=[-0.06,-0.02], $\mu_{T75r}$=[-0.61,-0.82]), *left* and 2-heptanone (OMP, [M-L,A-P]: $\mu_{\text{z-score amplitude}}$=[0.19,0.43], $\mu_{\text{odor-response corr}}$=[0.23,0.38], $\mu_{T75r}$=[-0.40,-0.51]; THY, [M-L,A-P]: $\mu_{\text{z-score amplitude}}$=[0.21,0.31], $\mu_{\text{odor-response corr}}$=[-0.03,0.14], $\mu_{T75r}$=[-0.51,-0.63]), *right*. (b) Histogram of the change in GI from odor intermittency 0.2 to odor intermittency 0.8 based on the linear regression fit of GI vs odor intermittency per glomerulus. (c) Average GI as a function of odor intermittency for sniff frequencies of 2-8 Hz (light to dark shades). (d) $GI_{nHz}$-$GI_{2Hz}$ per glomerulus for each intermittency value for methyl valerate, *top* and 2-heptanone, *bottom.* (e) Linear classifier performance (accuracy) over 240 glomeruli when trained on trials of four different sniff frequencies (2, 4, 6, 8 Hz) when presented with methyl valerate. 60 iterations (20 times 3-fold) per classifier. OMP: exponential plateau fit, 2 Hz: Y=0.91 -(0.34)*($e^{-0.03x}$), $r^2$=0.55; 8 Hz: Y=0.86 -(0.32)*($e^{-0.03x}$), $r^2$=0.42. THY: exponential plateau fit, 2 Hz: Y=0.92 -(0.5)*($e^{-0.05x}$), $r^2$=0.66; 8 Hz: Y=0.93 -(0.38)*($e^{-0.04x}$), $r^2$=0.61.

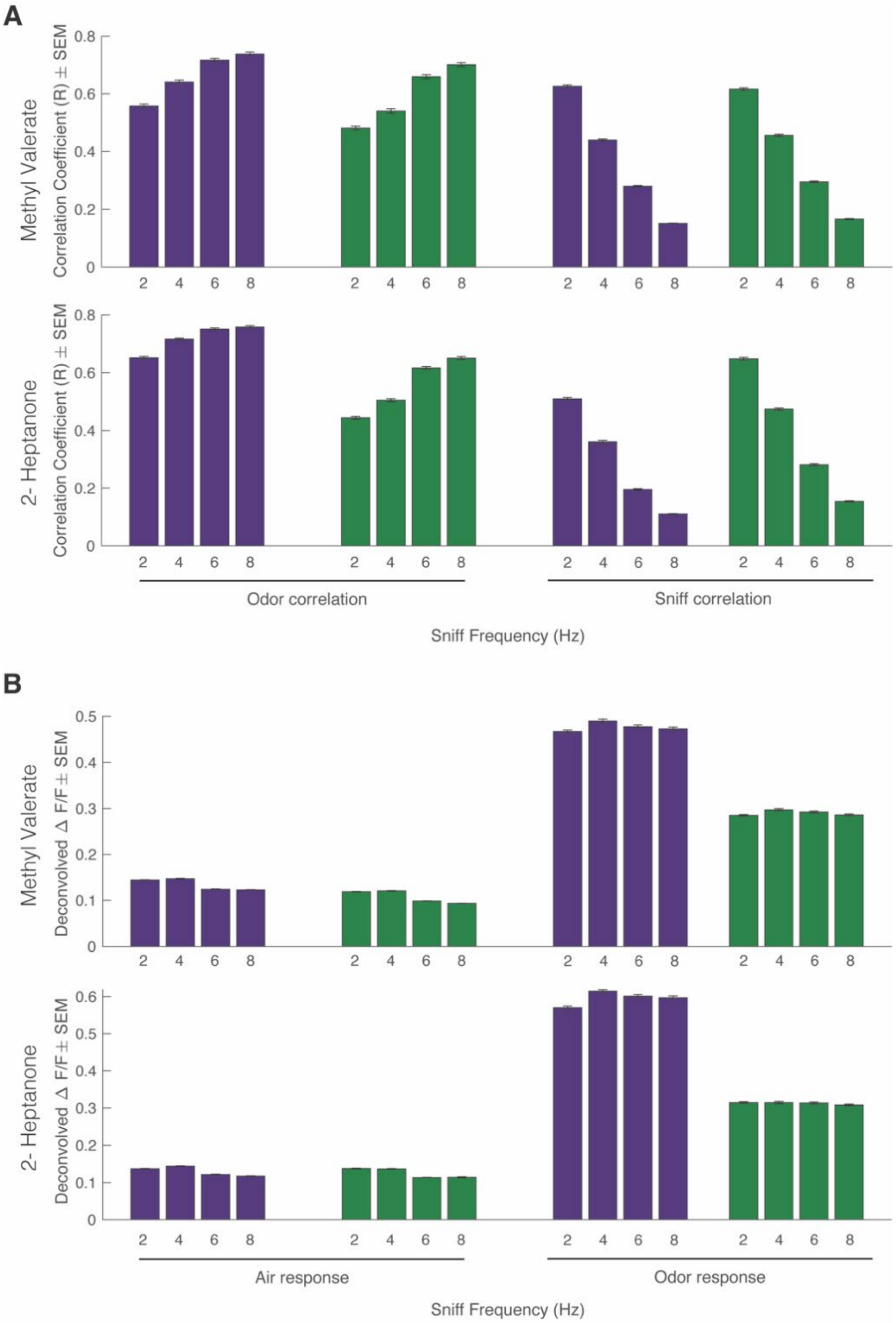


**Figure 5- figure supplement 2. Effect of sniff frequency on odor correlation, sniff correlation, air response, and odor response.** For all graphs purple indicates OSNs (OMP-GCaMP6f) and green indicates M/T cells (THY1-GCaMP6f). (a) *Top* Methyl Valerate, *Bottom* 2-Heptanone. *left* Average maximum cross-correlation between glomerular response and PID reading across sniff frequencies (Spearman correlation, $p<0.0001$). *right* Average maximum cross-correlation between glomerular response and pressure sensor reading (sniff measurement) across sniff frequencies (Spearman correlation, $p<0.0001$). (b) *Top* Methyl Valerate, *Bottom* 2-Heptanone. *left* Average glomerular deconvolved ΔF/F prior to odor presentation (Spearman correlation, $p<0.0001$). *right* Average glomerular deconvolved ΔF/F during odor presentation (Spearman correlation, $p>0.05$). (Methyl Valerate: OMP n= 367 glomeruli, 7 mice; THY n= 294 glomeruli, 6 mice; Heptanone: OMP n= 241 glomeruli, 6 mice; THY n= 271 glomeruli, 6 mice; n= 45 trials per sniff frequency, total n= 180 trials).
